# SEN1 is responsible for molybdate transport into nodule symbiosomes for nitrogen fixation in *Lotus japonicus*

**DOI:** 10.1101/2022.11.10.515970

**Authors:** Qingnan Chu, Tsuneo Hakoyama, Makoto Hayashi, Kiminori Toyooka, Mayuko Sato, Takehiro Kamiya, Toru Fujiwara

## Abstract

Symbiotic nitrogen fixation (SNF) in legume root nodules requires a steady supply of molybdenum (Mo) for synthesis of the iron-Mo cofactor for nitrogenase in bacteroids. For this nutrient to be exported by the host plant it must cross the peribacteroid membrane (PBM), however, the molybdate transporter responsible has not yet been identified. A *Lotus japonicus* symbiotic mutant, *sen1*, forms nodules that do not fix nitrogen; it has nodule defects and bacteroid degradation. The biochemical function and subcellular localization of SEN1 protein remains to be elucidated. Here, we found a new phenotype in which the *SEN1* mutation resulted in increased Mo accumulation in the nodule host fractions but decreased Mo accumulation in the bacteroids at 10 days post inoculation. We identified the molybdate efflux transport activity of SEN1 via heterologous expression in yeast. *SEN1* was expressed exclusively in nodules, and its expression was stable in response to varying Mo supply in nutrient solution. *In situ* immunostaining verified that the SEN1 protein is localized, in part, to the PBM in the rhizobium-infected cells. Taken together, these results confirmed that SEN1 is responsible for mediating molybdate efflux from the cytosol of nodule host cells to the symbiosomes for SNF. Furthermore, *SEN1* mutation reduced the expression of *nifD* and *nifK*, suggesting that *SEN1* may be pertinent to iron-Mo-cofactor assembly. This work fills the knowledge gap regarding how molybdate is allocated from the host plant to the bacteroids; such knowledge is critical for developing new SNF biological systems in non-legume plants.

**One-sentence summary:** SEN1 is localized partly in the peribacteroid membrane of nodule cells and mediates the molybdate exportation from the host plant cytosol to the symbiosomes for symbiotic nitrogen fixation.

## Introduction

Symbiotic nitrogen fixation (SNF) performed in differentiated root organs (nodules) by the legume–rhizobia partnership is one of the main alternatives to the overuse of synthetic N fertilizers (Henneron et al., 2020; Herridge et al., 2022). Within nodules, endosymbiotic rhizobia differentiate into bacteroids surrounded by a peribacteroid membrane (PBM) (Day et al., 1989), forming the organelle-like structures—symbiosomes—where SNF takes place. The enzyme that catalyzes SNF is a metalloprotein called a nitrogenase—the only one in the biosphere that converts N2 into ammonium (Milton, 2022). In exchange for this ammonium, the host plant provides photosynthates and mineral nutrients for the bacteroids. Nutrient exchange between the host plant and the bacteroids across the PBM is essential for SNF (White et al., 2007; Udvardi and Poole, 2013).

Molybdenum (Mo) is one of the most crucial nutrients transferred to the bacteroids, because it is required for assembly of the iron (Fe)-Mo cofactor (FeMoco) of nitrogenase (González-Guerrero et al., 2014; Ohki et al., 2022). Additionally, Mo is essential for plants as a part of the Mo cofactor in five enzymes involved in nitrate assimilation, phytohormone biosynthesis, purine metabolism, and sulfite detoxification (Hille et al., 2011). Mo, unlike other transition metals, is taken up from the soil as the oxyanion molybdate (MoO_4_^2-^) instead of in a cationic form. To satisfy the symbiotic requirement and supply the bacteroids, these MoO_4_^2-^ must first be translocated to the rhizobia-infected cells of the nodules and then transverse the PBM and the space between the PBM and the bacteroid membrane (Tejada-Jiménez et al., 2017; Gil-Díez et al., 2019). The *ModABC* operon is responsible for molybdate uptake from the peribacteroid space, which is followed by Mo incorporation into the bacteroids (Hernandez et al., 2009). The only known plant-type-specific molybdate transporters belong to the Molybdate Transporter type (MOT) family; the first one was identified in the higher plant *Arabidopsis thaliana* (Tomatsu et al., 2007). In legume plants, *LjMOT1* has been identified to play a role in molybdate uptake from the soil and translocation to the shoots of *Lotus japonicus* (Gao et al., 2016; Duan et al., 2017). *MtMOT1.2* has been identified in *Medicago truncatula* to mediate molybdate delivery via the vasculature into the nodules (Gil-Díez et al., 2019), and *MtMOT1.3* further introduces molybdate into rhizobia-infected or uninfected cells within the nodules (Tejada-Jiménez et al., 2017). However, the transporters mediating Mo loading from nodule host cells to the symbiosomes for SNF remain unknown.

The mutant *sen1 (<u>s</u>tationary <u>e</u>ndosymbiont <u>n</u>odule 1*) forms ineffective nodules (Fix^-^phenotype), blocks N fixation, and impairs the bacteroid differentiation in *L. japonicus* (Suganuma et al., 2003; Hakoyama et al., 2012). Expression of *SEN1* has been detected exclusively in the nodules, and Southern blot analyses have revealed that the *SEN1* clade appears to be specific to legumes (Hakoyama et al., 2012). *SEN1* encodes an integral membrane protein homologous to CCC1 (Ca^2+^-sensitive cross-complementer 1), which is a vacuolar Fe/manganese transporter in *Saccharomyces cerevisiae*, and VIT (vacuolar iron transporter) in *A. thaliana* (Liu et al., 2020). The orthologous gene in soybean, *GmVTL1a (Glycine max vacuolar iron transporter-like*), is responsible for Fe transport across the PBM to bacteroids (Brear et al., 2020; Liu et al., 2020). However, the yeast complementation assays have demonstrated that *SEN1* cannot complement the *Δccc1* mutant in yeast (Brear et al., 2020), suggesting that it may not be responsible for Fe transport. Although SEN1 protein is essential to the N fixation capacity of legume nodules, the physiological role and subcellular localization of this protein remains to be elucidated.

Here, on the basis of the results of a phenotypic analysis and the heterologous expression of *SEN1* in *S. cerevisiae*, we demonstrated that SEN1 is a molybdate efflux transporter. Also, we demonstrated that SEN1 protein is localized, in part, to the PBM, thus allowing it to mediate the molybdate transport from the host plant cytosol to the symbiosome by crossing the PBM. This is the first molybdate efflux transporter known so far, and out findings answer the question of how molybdate is transported from the host cell cytosol to the bacteroids in the nodules.

## Results

### *sen1* mutants show defects in nodule development and plant growth under N deficiency

*sen1-1* and *sen1-2* mutants were produced from ethylmethane sulfonate metagenesis of *L. japonicus* ecotype Gifu B-129. Both harbored single nucleotide mutations (*sen1-1*, C122T; *sen1-2*, G332A) leading to amino acid substitutions (*sen1-1*, A41V; *sen1-2*, R111K) (**Figure S1**), as reported in previous studies (Kawaguchi et al., 2002; Hakoyama et al., 2012). *sen1-1* and *sen1-2* formed nodules upon inoculation with *Mesorhizobium loti* under N-free conditions. However, they displayed N-deficiency symptoms under symbiotic conditions (**Figure 1A**), including yellow leaves, a concomitant increase in the root to shoot ratio, and anthocyanin production in the stem. In *sen1* mutants these phenotypes have been attributed to a deficiency of N fixation (Suganuma et al., 2003). The nodules that formed on *sen1-1* and *sen1-2* were obviously smaller than those on the wild type (WT) *Gifu* (**Figure 1B**). Pink is a mark of nodule maturity because of the presence of abundant leghemoglobin, as occurred in the 78.5% of nodules on the WT; in contrast, the nodules formed on *sen1-1* and *sen1-2* were almost white, with only 5.4% and 10.6% of them, respectively, pink (**Figure 1C**).

**Figure 1.**
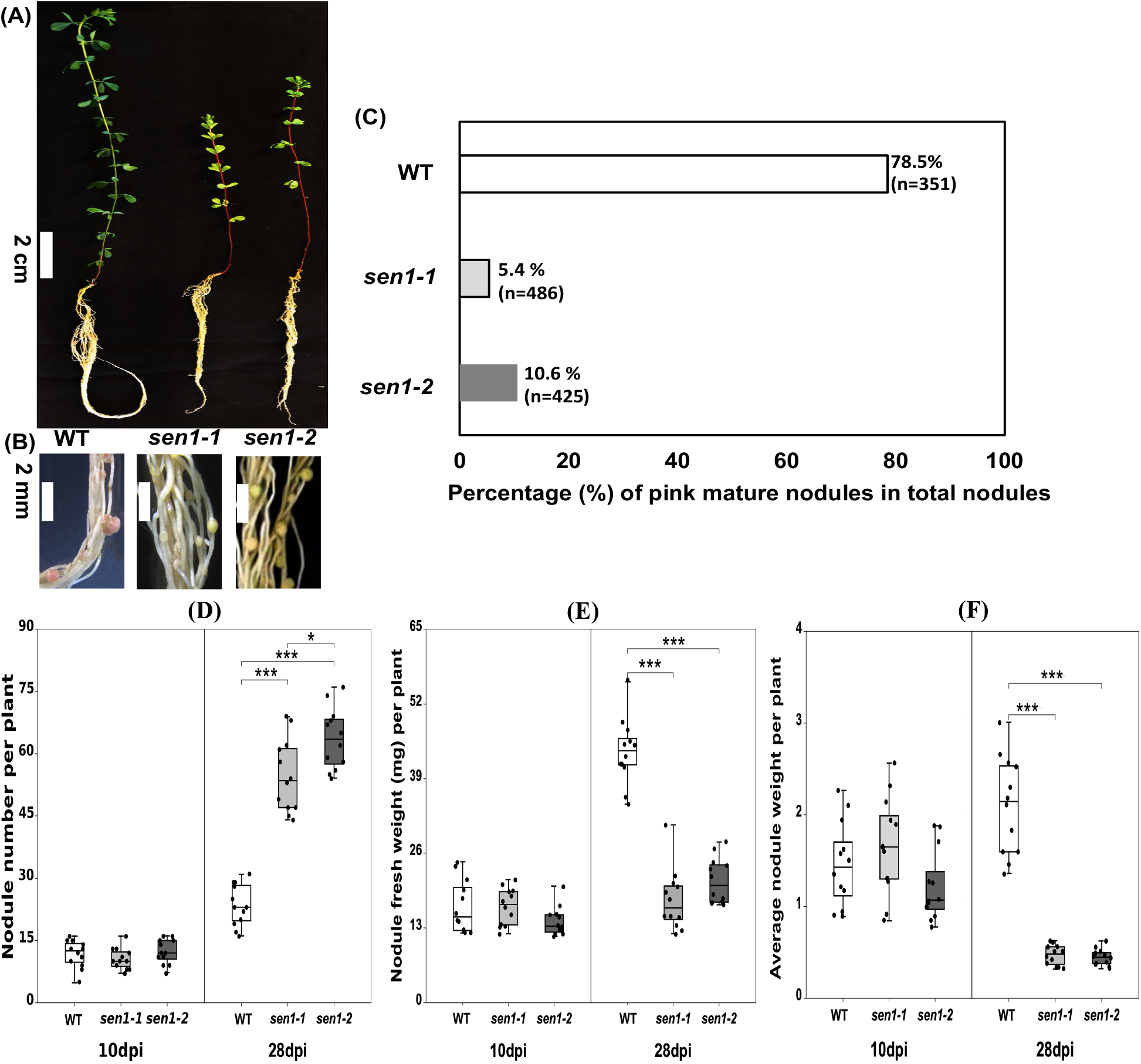
Phenotypic analysis of *sen1-1* and *sen1-2* mutant plants after rhizobial inoculation. (A) Growth of *Lotus japonicus* in the pots filled with vermiculites and watered with B&D medium (Mo concentration: 170 nM). Scale bar = 2 cm. (B) Nodule growth. Scale bar = 2 mm. (C) Percentage of mature pink nodules in total nodules at 28 dpi. Scale bar = 1mm. (D) Nodule number per plant. (E) Nodule fresh weight per plant. (F) Average nodule weight. Data are means ± standard deviation (Student’s *t* test, two-tailed *P* < 0.05 *, *P* < 0.01 **, *P* < 0.001 ***, n = 12). dpi: days past inoculation.

Nodule number, total fresh weight and average weight of nodules per plant were quantified at 10 and 28 days post-inoculation (dpi), respectively. At 10 dpi inhibition of bacteroid differentiation has been observed in *sen1* mutants (Hakoyama et al., 2012). At 28 dpi WT nodules generally reach maturity (Tobergte and Curtis, 2013). At 10 dpi no significant differences were observed in nodule number or total fresh weight or average weight of nodules per plant between the mutants and the WT (**Figure 1D–F**). At 28 dpi the nodule numbers in *sen1-1* and *sen1-2* were 2.3 and 2.8 times higher than that in WT (both *P* < 0.001), whereas the total nodule fresh weight and average nodule weight per plant were significantly lower in both mutants (both *P* < 0.001) (**Figure 1D–F**). These observations indicated that the *sen1* mutation resulted in plant growth defects under N fixation deficiency caused by altered nodulation in *L. japonicus*.

### *SEN1* mutation alters Mo allocation from the host plant fraction to bacteroids in nodules

Bacteroids were separated from the host-plant fraction of the nodule cells. At 10 dpi, Mo accumulation in the whole nodules was comparable in the *sen1* mutants and the WT, irrespective of whether the Mo supply was sufficient or deficient (**Figure 2A** and **2D**). However, at 10 dpi, with sufficient Mo supply (0.17 μM Na_2_MoO_4_), the Mo concentration in the host plant fraction in *sen1-1* and *sen1-2* was significantly higher than that in the WT whereas in bacteroids it was significantly lower (both *P* < 0.001) (**Figure 2B** and **2C**). At 10 dpi, with deficient Mo supply, although the Mo concentration in the host plant fraction was comparable between the *sen1* mutants and the WT, in the bacteroids of *sen1-1* and *sen1-2* it was significantly lower than in the WT (both *P* < 0.001) (**Figure 2E** and **2F**). At 28 dpi, a trend of significantly higher Mo accumulation in the nodules of the WT was observed in the whole nodules, in the host plant fraction, and in the bacteroids under either Mo sufficiency or deficiency (all *P* < 0.001) (**Figure 2**). This can be ascribed to reduced Mo importation to the nodules because of the inhibition of nitrogenase activity in the *sen1* mutants (Suganuma et al., 2003). Furthermore, we observed no significant differences in the Mo concentration in the roots or shoots between *sen1* mutants and the WT, regardless of the growth duration or Mo supply (**Figure S2**). These results suggest that, at 10 dpi, the *SEN1* mutation alters Mo allocation between the host plant host fraction and the bacteroids, possibly inhibiting Mo supply from the host plant to the bacteroids.

**Figure 2.**
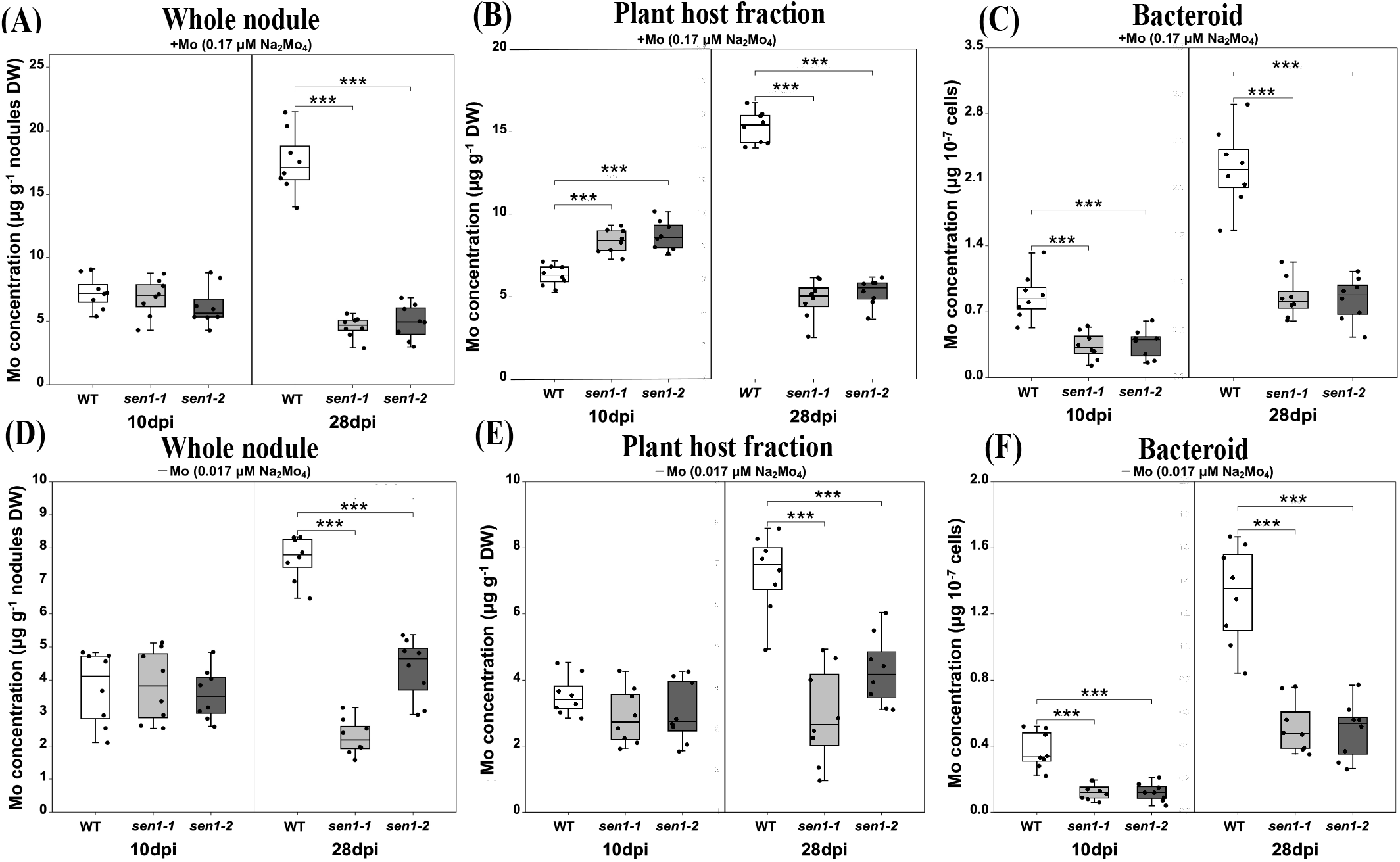
Effects of *SEN1* knockdown on Mo distribution in nodules of *Lotus japonicus* under sufficient (ABC) and deficient (DEF) Mo supply. (AD) Mo concentration in the whole nodules; (BE) Mo concentration in the plant host fraction; (CF) Mo concentration in the bacteroids. Data are means ± standard deviation of two independent experiments and each one contains 4 replicates. (Student’s *t* test, two-tailed *P* < 0.05 *, *P* < 0.01 **, *P* < 0.001 ***). dpi: days past inoculation.

### Heterologous expression of *SEN1* in *S. cerevisiae* shows Mo efflux transport activity

*SEN1* consists of only one exon without an intron, encoding a peptide with 227 amino acids. Domain prediction suggested that SEN1 is an integral membrane protein with five transmembrane regions (**Figure S1**). The altered molybdate allocation between the plant host fraction and the bacteroids in *sen1* mutants (**Figure 2**) suggested that *SEN1* was involved in molybdate transport. Therefore, the *SEN1* and *sen1-1* genes were introduced into *S. cerevisiae* (BY4741) by using the pYES2 expression vector, which allows the expression of the inserted gene under the control of the galactose-inducible *GAL1* promoter. After being grown in SD (with glucose as a carbon source) or SG (with galactose as a carbon source) medium to the mid-log phase, the transformed yeast cells were transferred to SD or SG medium containing 170 nM molybdate for 30 min, after which the element concentrations in the cells were determined (**Figure 3**). When the yeast cells were grown with glucose, Mo accumulation in the cells was equally low among the different transformants. When the glucose in the medium was replaced with galactose, the Mo concentration in the yeast cells carrying *SEN1* was 5.4 times lower than that in the cells carrying the empty vector and 5.1 times lower than that in the *sen1-1* cells (A41V) (*P* < 0.001) (**Figure 3A**). In contrast, the yeast cells carrying *SEN1*, after being grown in SG medium, accumulated contents of sulfur (S), Fe, Mn, copper (Cu), and zinc (Zn) similar to those in the yeast cells carrying empty vector (**Figure 3B**). Fe, Mn, Cu, and Zn were analyzed because *SEN1* is orthologous to *CCC1* and *VIT* (Hakoyama et al., 2012). S was analyzed because molybdate and sulfate are chemically similar and the first molybdate transporter in plants, MOT1, was originally annotated as a sulfate transporter (Sultr5;2) (Tomatsu et al., 2007). These results indicated that, despite its being orthologous to De or Mn transporters, SEN1 owns the capacity for Mo efflux transport, and the point mutation in *sen1-1* (A41V) resulted in the loss of Mo efflux activity.

**Figure 3.**
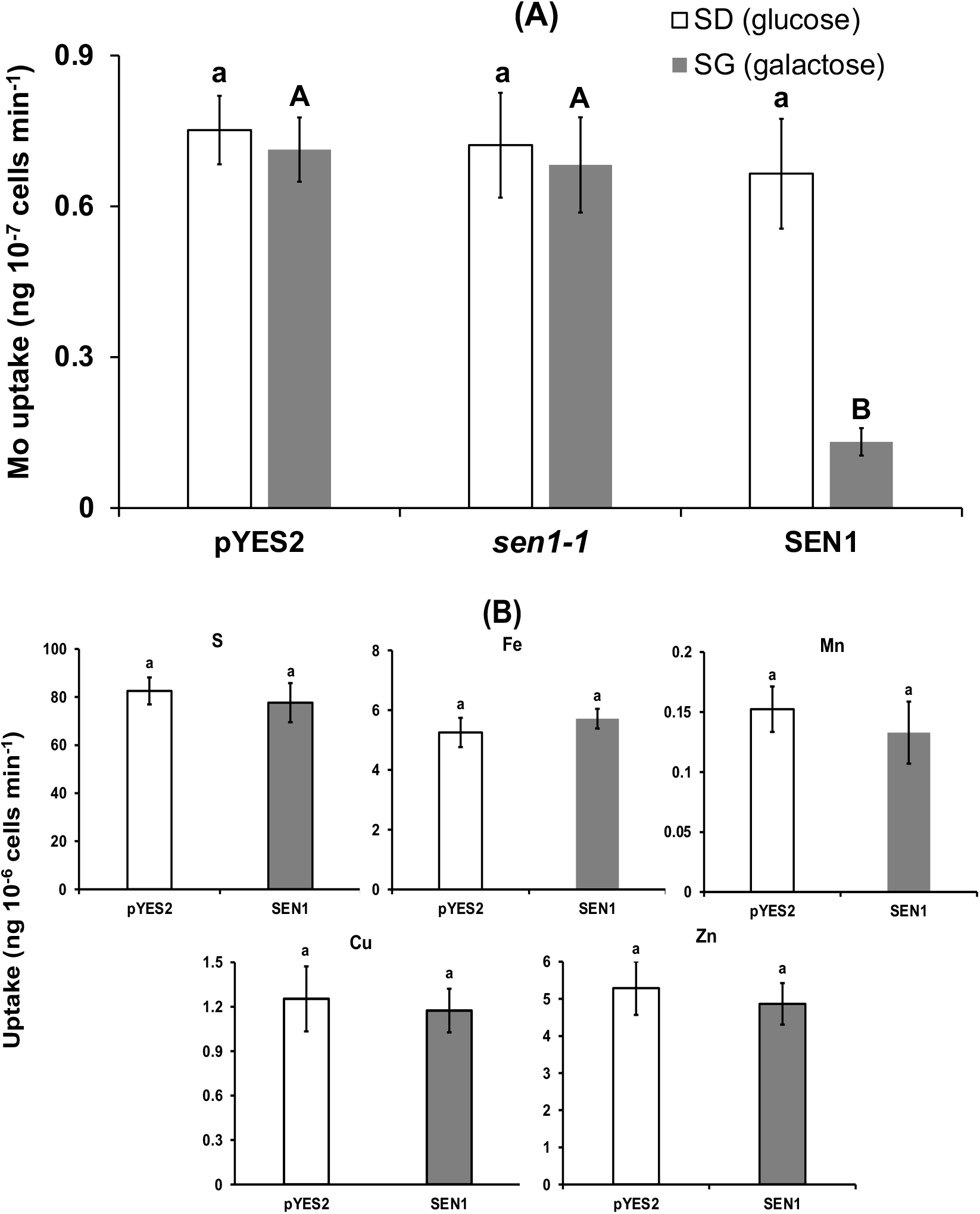
Mo transport activity of *SEN1* in *Saccharomyces cerevisiae*. (A) Mo uptake in yeast cells expressing *SEN1* containing vector control (pYES2) or cloned from wild type (*Gifu*) or *sen1-1*. Yeast cells were incubated in selective media containing glucose (SD) or galactose (SG) supplemented with 170 nM MoO_4_^2-^ for 30 min. Different letters indicate significant differences (Duncan’s test, *P* < 0.001, n = 3 in each independent experiment for two times); small letters a-b; glucose; capital letters A-B: galactose. (B) Sulfur (S), iron (Fe), manganese (Mn), copper (Cu), and zinc (Zn) concentration in the yeast cells expressing *SEN1* or containing vector control (pYES2). Yeast cells were incubated in selective media containing galactose. Data are means ± standard deviation.

### *sen1* mutants downregulate the expression of *nifD* and *nifK*

*nifD* and *nifK* encode the α-subunit and *β*-subunit, respectively, of the heterotetrameric FeMo protein of nitrogenase. This protein is essential for the incorporation of Mo during nitrogenase biosynthesis. Expression of *nifD* and *nifK* in the bacteroids of the *sen1* mutants and the WT was detected by quantitative RT-PCR analysis (**Figure 4**). The expression levels of *nifD* in the bacteroids of *sen1-1* and *sen1-2* were, 4.5 and 3.3 times lower than that of WT, respectively, and those of *nifK* were 3.5 and 2.7 times lower, respectively. These results indicated that mutation in *SEN1* had negative impacts on the assembly of FeMoco.

**Figure 4.**
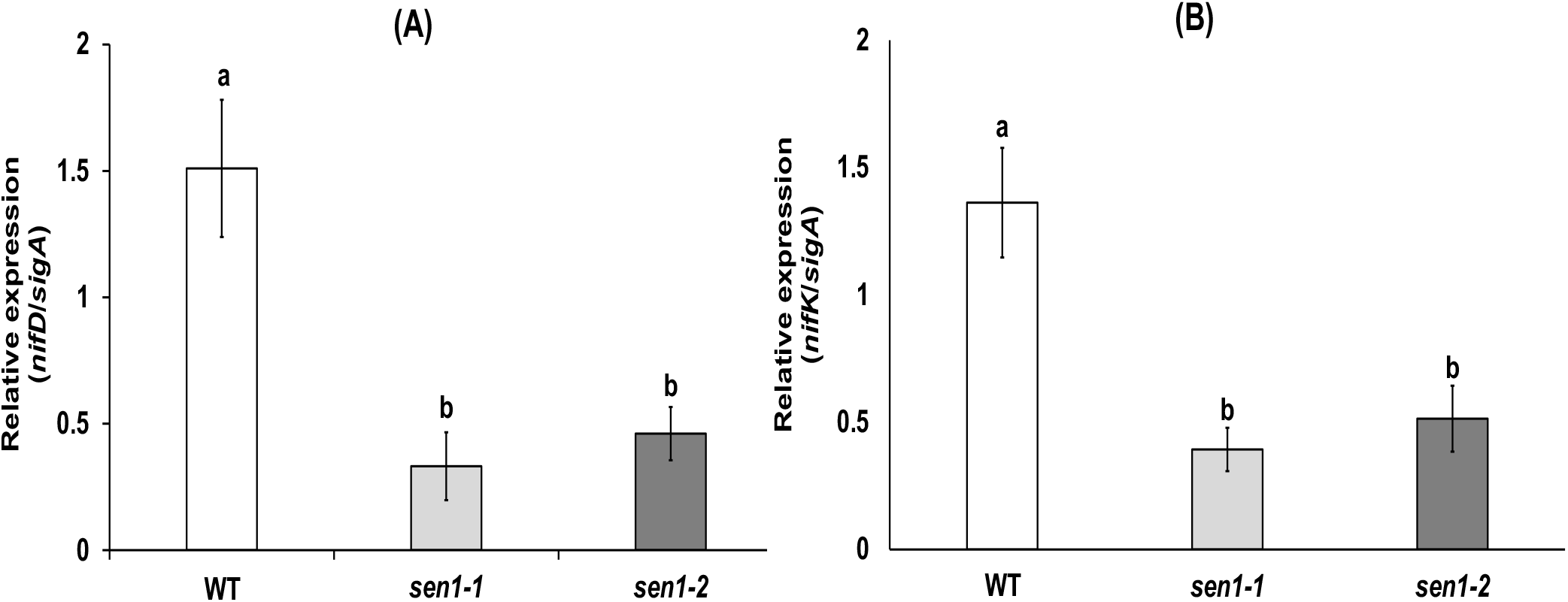
Expression of *nifD* (A) and *nifK* (B) in nodules formed on the wild-type plant and the *sen1-1* and *sen1-2* mutants at 28 dpi. Expression was evaluated with qRT-PCR using *sigA* as an internal standard. Data are means ± standard deviation (Duncan’s test, *P* < 0.001, n = 3).

### SEN1 is, in part, localized to the PBM of nodule cells

A previous study has demonstrated that *SEN1* is expressed exclusively in the nodules (Hakoyama et al., 2012). We confirmed this by measuring the *SEN1* transcript abundance in different organs under different levels of Mo supply at 28 dpi by using quantitative RT-PCR (qRT-PCT) (**Figure 5A**). The results agreed well with those of the abovementioned study and showed that *SEN1* transcript was abundant in the nodules but almost undetectable in the other plant organs examined. Additionally, the expression level of *SEN1* was insensitive to variations in Mo concentration in the nutrient solution (**Figure 5A**). Moreover, we investigated the temporal dynamics of *SEN1* expression in the nodules (**Figure 5B**). Expression of *SEN1* was relatively low in young nodules (7 dpi). It then increased gradually, peaked at 28 dpi, and then steadily declined in older nodules. This may be attributable to reduced nutrient supply to nodules after 28 dpi because of inhibited nodule development. This result is consistent with a previous Northern blot analysis of temporal SEN1 expression in the nodules (Hakoyama et al., 2012).

**Figure 5.**
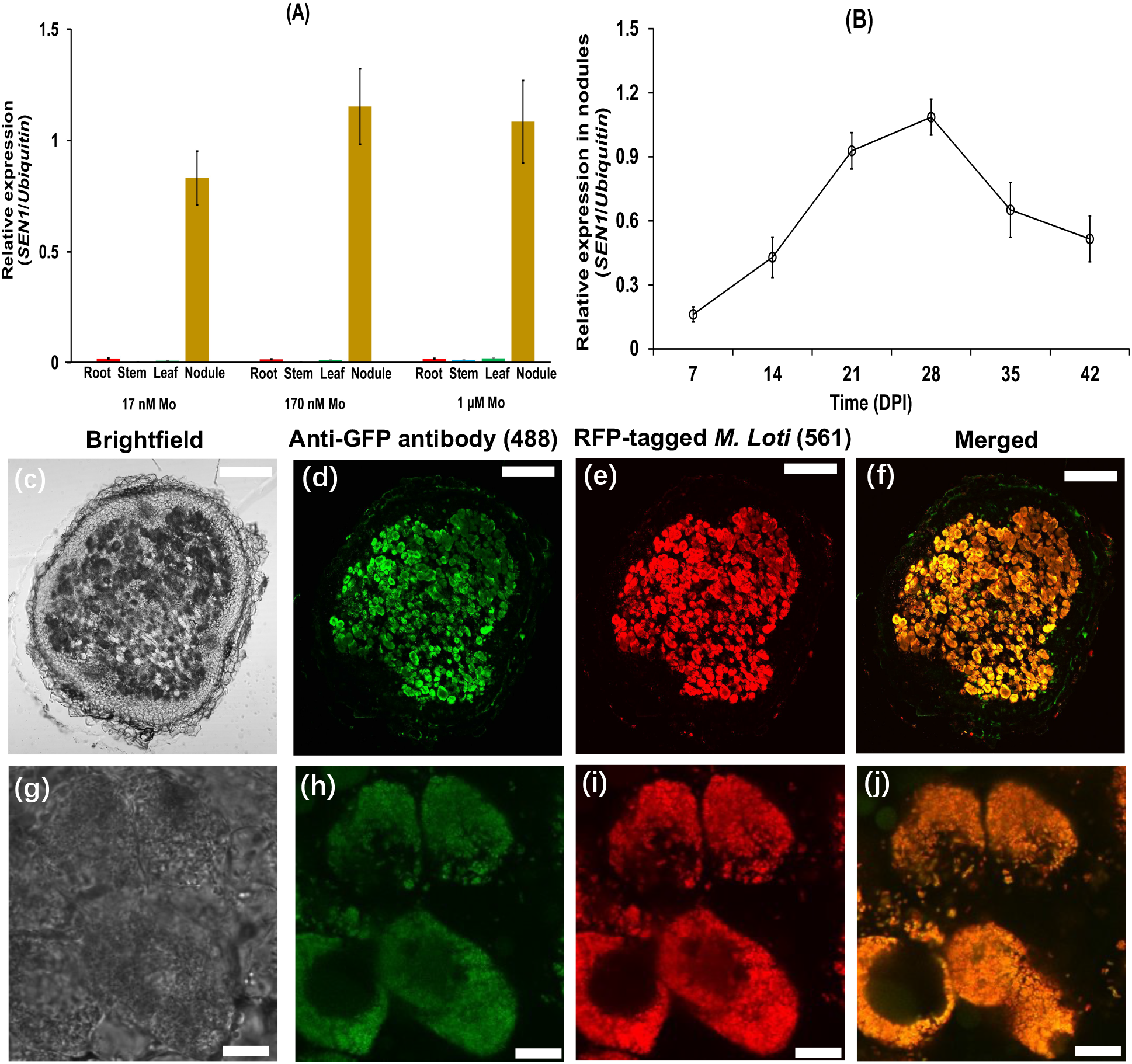
Tissue specificity of *SEN1* expression and subcellular localization of SEN1 protein in *Lotus japonicus* nodules. (a) Transcript level of *SEN1* in tissue samples from roots, stems, leaves and nodules under different level of Mo supply at 28 dpi (n = 3). (b) Time-course transcript level of *SEN1* during nodule development. Expression was evaluated by qRT-PCR using ubiquitin as an internal standard. Data are means ± standard deviation (n = 3). (c–j): Immunofluorescence staining of*pSEN1:SEN1-GFP* transgenic nodules in hairy roots. The infected zone in (c–f) are magnified in (g–j). Hairy roots carrying *pSEN1:SEN1-GFP* were inoculated with a *M. Loti* strain carrying an RFP plasmid. 21-dpi nodule cross-sections were immunostained with a specific antibody against GFP. Green shows anti-GFP signals with excitation wavelength at 488 nm. Red shows RFP-tagged rhizobia with excitation wavelength of 561 nm. Bars: (c-f) 200 μm; (g-j) 10 μm.

To elucidate the cellular and subcellular localization of SEN1, we used hairy root transformation to introduce an N-terminal-green fluorescent protein (GFP) fusion with the *SEN1* coding sequence, under the control of the native promoter of *SEN1. In situ* immunostaining of the *pSEN1:SEN1-GFP* product was conducted in transgenic nodules from hairy roots inoculated with red fluorescence protein (RFP)-expressing *M. loti* MAFF303099. The observed anti-GFP antibody signals targeted mainly those cells that were colonized by rhizobia (**Figure 5C–J**), implying that SEN1 functions in rhizobium-infected cells. Furthermore, the anti-GFP signals from the infected cells overlapped well with the RFP signals from the rhizobia (**Figure 5F and 5J**), suggesting that SEN1 was localized to the symbiosomes.

To further clarify the subcellular localization of SEN1, we performed immunolocalization of SEN1 using immunoelectron microscopy. The antibody was obtained by immunizing rabbits with a synthetic peptide (positions 111 to 128 of the SEN1 amino acid sequence). The specificity of the antibody against SEN1 protein was confirmed by western blot using the total and the microsome fractions of nodules from the WT (**Figure S3**). A single band was observed for both fractions, and the size seemed to accord with the predicted SEN1 protein size (23,774 Da). With this antibody, SEN1 localization was further determined by using transmission electron microscopy and a colloidal-gold-conjugated secondary antibody (**Figure S4**). In the cross-sections of WT nodule samples obtained at 7, 14, and 21 dpi, the epitope was detected at the PBM, in bacteroids, and in the intracellular components of host nodule cells, including the cytosol and endomembrane compartments. The frequency of colloidal gold particles observed at the PBM in nodule samples was 35.6% at 7 dpi (n = 118), 42,1% at 14 dpi (n = 159), and 40.4% at 21 dpi (n = 75) (**Figure S5**). These results indicated that SEN1 was localized, in part, at the PBM of rhizobium-infected cells in nodules.

## Discussion

Acquisition of N from the atmosphere by SNF is a sustainable alternative to the intensive use of synthetic N fertilizers in agriculture (Henneron et al., 2020; Herridge et al., 2022). Mo supply is critical for SNF, as this micronutrient is essential for synthesis of the enzyme nitrogenase, which is directly involved in the reduction of N2 to the phytoavailable N source, ammonium. However, in spite of the importance of Mo in SNF, the proteins mediating molybdate loading to symbionts remains to be discovered. Once in the cytosol of plant host cells, molybdate has to be pumped out and transported across the PBM to engage the bacteroids in nitrogenase synthesis. However, to our knowledge, no molybdate-specific efflux system is known. The only known plant-type specific molybdate transporters are all from the MOT family and show influx activity, including *MOT1* and *MOT2* in Arabidopsis (Tomatsu et al., 2007; Baxter et al., 2008; Gasber et al., 2011), *OsMOT1;1* and *OsMOT1;2* in rice (Huang et al., 2019; Ishikawa et al., 2021; Hu et al., 2022), *MtMOT1.2* and *MtMOT1.3* in *Medicago Truncatula* (Tejada-Jiménez et al., 2017; Gil-Díez et al., 2019), and *LjMOT1* in *L. japonicus* (Gao et al., 2016; Duan et al., 2017). Considering the renewed interest in introducing N2-fixing capacity into nonlegume cereals and the need for sustainable agriculture (López-Torrejón et al., 2016; Zhao et al., 2022), this knowledge gap needs to be filled, with the aim of ensuring Mo delivery to produce functional nitrogenase in new biological systems. Our results are a decisive step towards the optimization of molybdate allocation for N_2_ fixation.

Here, we have identified a nodule-specific molybdate efflux transporter, from a new family, as responsible for efficient Mo supply from the cytosol of host plant cells to bacteroids for SNF.

*sen1* belongs to the Fix-class of mutants, which form nodules filled with endosymbionts with dramatically reduced SNF activity, defects in bacteroid differentiation, and impaired nodule development (Kawaguchi et al., 2002; Suganuma et al., 2003), implying that *SEN1* is required for symbiosome maturation and SNF. These characteristics have also been discovered in the phenotype of *Gmvtl1* mutants (Brear et al., 2020; Liu et al., 2020). *GmVTL1a* in soybean is an ortholog of *SEN1* but encodes a ferrous (Fe^2+^) transporter facilitating Fe^2+^ import into the vacuole when expressed in yeast. However, yeast complementation assay results have shown that *SEN1* cannot rescue the *Δccc1* mutant yeast, although *GmVTL1a* could (Brear et al., 2020). Our findings demonstrated that the heterologous expression of *SEN1* in yeast showed molybdate efflux capacity. Although Brear et al., (2020) provided indirect envidence for the potential role of *SEN1* in Fe transport (rhizonium-infected cells in WT nodules showed stronger Perls/diaminobenzidine-staining than infected cells in *sen1-1* nodules), molybdate deficiency in symbionts may result in lower Fe accumulation, because Mo and Fe are both required by the bacterods, and it is possible that their respective transporters are possibly co-expressed. *SEN1* and *GmVTL1a* have 66% amino acid identity, whereas both of the protein structures of both remain unclear. Our previous study also demonstrated that the *SEN1* mutation resulted in reduced Fe concentration in both the host plant fraction and the bacteroids (Hakoyama et al., 2012). This reduced Fe accumulation in the infected cells of *sen1* nodules may be attributed to a lower Fe requirement because Mo shortage inhibits nitrogenase biosynthesis, rather than to a loss of PBM-localized Fe transporter. The different functions of the orthologous genes *SEN1* and *GmVTL1a* may be associated with the differences in the unknown domain (**Figure S1**). Further investigation of the protein structure is needed to elucidate the differences in function between *SEN1* and *GmVTL1a*.

We found here that increasing the molybdate supply in the nutritive solution had a negligible impact on *SEN1* expression in the nodules (**Figure 5A**) or on Mo accumulation in the bacteroids of *sen1* mutants (**Figure 2**). Additionally, Suganuma et al., (2003) have reported that mutation of *SEN1* completely abolishes SNF capacity. These results suggest that other genes are unable to complement the loss of *SEN1* function in *sen1* mutants. This is different from the case with *MtMot1.2* and *MtMOT1.3*, two other nodule-specific Mo transporters that have been characterized (Tejada-Jiménez et al., 2017; Gil-Díez et al., 2019): 25%and 12% nitrogenase activity remained in the *mot1-2* and *mot1-3* nodules, respectively, and the phenotype could be rescued by increasing the molybdate supply (Tejada-Jiménez et al., 2017; Gil-Díez et al., 2019). This implies that, at high molybdate concentration, other membrane transporters could counterbalance the absence of *MtMOT1.2* or *MtMOT1.3* activity. Although there are candidate genes, namely homologs of *LjMOT1* (Duan et al., 2017) or *SEN1*, which are predicted to be expressed in a nodule-specific manner, seemingly none of them enables the compensation of *SEN1* mutation. This implies that *SEN1* is ultimately more important for SNF than it is for Mo nutrition, because it is involved in SNF and is expressed exclusively in nodules. Therefore, further studies are needed to investigate whether *SEN1* is the sole transporter for the supply of molybdate into symbionts. Interestingly, phenotypes similar to those of *sen1* has been reported in the loss-of-function of *SST1* (PBM-localized *symbiotic sulfate transporter* in *L. japonicus*) (Krusell et al., 2005) and *GmVTL1a* (Brear et al., 2020; Liu et al., 2020). In theory, the reduced Fe supply resulting from *GmVTL1a* knockout can be complemented by the action of other known PBM-localized Fe exporters, including *DMT1 (divalent metal transporter 1*) (Kaiser et al., 2003) and *FPN2 (ferropotin2*) (Escudero et al., 2020). However, the complementation does not occur, even during growth under Fe sufficiency, and the *GmVTL1a* mutant has nodule defect identical to those in *sen1* (Liu et al., 2020).

*SEN1* is expressed exclusively in the nodules (**Figure 5A**), in agreement with previous reports that genomic Southern hybridization does not detect DNA fragments homologous to *SEN1* in non-legume plants (Hakoyama et al., 2012). *SEN1* expression in the nodules increased continuously from 7 to 28 dpi (**Figure 5B**). This expression profile is consistent with a situation in which the rhizobia-infected cells are increasing their molybdate content to supply it to the bacteroids and thus satisfy the increasing demand for FeMoco synthesis as the nodule develops. After 28 dpi, the Mo requirement of the bacteroids might become lower, thus leading to a decline in *SEN1* expression (**Figure 5B**).

*In situ* immunostaining has shown that the antibody signal from native promoter-driven SEN1-GFP is localized at symbiosomes. Although it was difficult to observe signals at the PBM due to limited resolution, we speculate that SEN1 is a PBM-localized protein, on the basis of the following evidence. First, the primary role of Mo in SNF is to be involved in the assembly of FeMoco for the synthesis of FeMo protein. *nifD* and *nifK* encode the α-subunit and *β*-subunit of the heterotetrameric FeMo protein, respectively. The expression levels of *nifD* and *nifK* in the bacteroids of *sen1* mutants were much lower than that in WT plants (**Figure 4**), suggesting that *SEN1* may be pertinent to the synthesis of the FeMo protein (**Figure 4**). Secondly, the molybdate content of the isolated bacteroids of *sen1* mutants was significantly lower than that of WT plants (**Figure 2**), suggesting that *SEN1* facilitates molybdate export from the cytosol of host plant cells into the symbiosomes. Finally, most components of the symbiosomes are derived from rhizobia, except for the PBM, which originats in the host plant. The overlapping signals between anti-GFP antibody and RFP-tagged rhizobium imply that the *L. japonicus* genome-encoded *SEN1* probably targets the PBM. Additionally, our immunolocalization using anti-SEN1 antibody showed that the signals were observed not only at the PBM, but also in the bacteroids, as well as in the cytosol and endomembrane of host cells. The epitope detected in the intracellular compartment, especially the membrane compartments, could correspond to newly synthesized SEN1 protein being ferried towards the PBM. The epitope detected in the bacteroids might represent non-specific signals against bacteroid components, such as peptidoglycan, or the signals might be indicative of some unknown function of SEN1 associated with the bacteroids. Taken together, these findings indicate that SEN1 is, in part, localized at the PBM in rhizobium-infected cells.

Another interesting finding presented here was that the *sen1* mutation resulted in the formation of more nodules (**Figure 1D**), consistent with the findings of a previous study (Suganuma et al., 2003). A similar phenotype has been discovered in the mutants of PBM-localized Fe or sulfate transporters, including *DMT1* (Kaiser et al., 2003), *GmVTIL1a* (Brear et al., 2020; Liu et al., 2020), *MtVTL4* (Walton et al., 2020), and *SST1* (Krusell et al., 2005). Notably, Fe, S, and Mo are all required for the synthesis of nitrogenase (González-Guerrero et al., 2014). Considering that transporter genes are usually downstream targets in regulatory networks, *SEN1* is unlikely to regulate autoregulation of a nodulation signaling pathway. One possible mechanism by which such pathways are regulated is by the feedback effect of inhibited SNF activity and nodule development, leading to the magnification of N-deficiency-responding signals and subsequently the stimulation of the generation of new nodules.

In this study, we demonstrated that *SEN1* encodes a molybdate efflux transporter and is required to export molybdate from the cytosol of host plant cells to the bacteroids. We also found that *SEN1* is exclusively expressed in the nodules of *L. japonicus* and is localized partly at the PBM. Because a steady supply of Mo is required for the synthesis of nitrogenase in the bacteroids of root nodules, *SEN1* is indispensable for SNF. We elucidated why *sen1* mutants showed inhibition of nitrogenase activity and nodule development, as reported in previous studies (Suganuma et al., 2003; Hakoyama et al., 2012). We propose a model describing the potential role of *SEN1* in molybdate transport for SNF in the nodule cells of *L. japonicus* (**Figure 6)**. SEN1 localizes at the PBM to mediate molybdate efflux from the cytosol of host plant cells to the symbiosome. When the molybdate reaches the peribacteroid space, *ModABC* (encoding a molybdate transporter of the ATP-binding cassette protein) introduces the molybdate into the bacteroids. These molybdate is loaded for the assembly of FeMoco and further for the biosynthesis of nitrogenase. Our results fill the knowledge gap regarding how molybdate is allocated from the host plant to the bacteroids for SNF. This information is critical for developing new SNF biological systems in non-legume plants.

**Figure 6.**
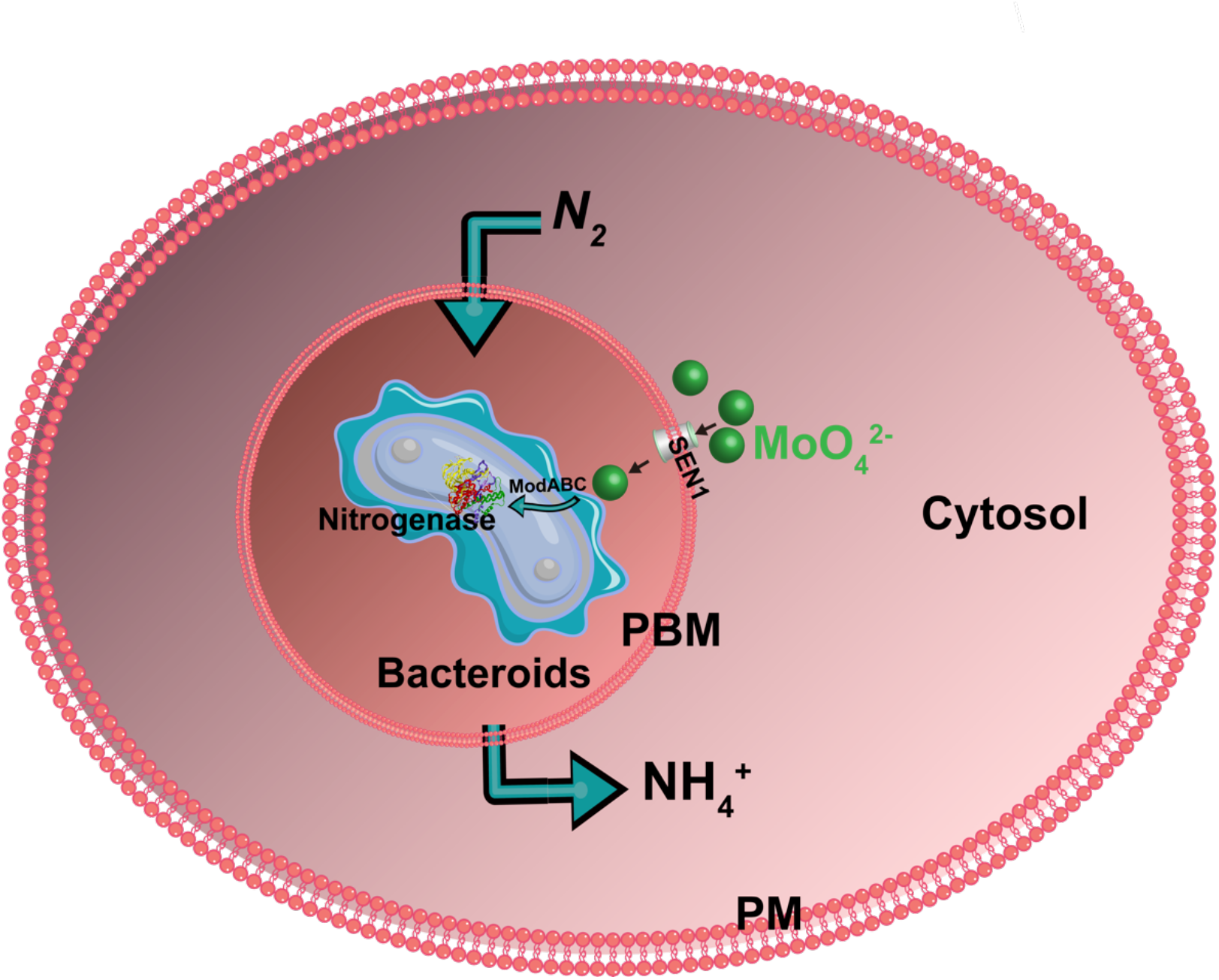
Proposed model of the potential role of *SEN1* in molybdate transport for nitrogen fixation in the nodule cells of *Lotus japonicus*.*SEN1* localizes on the peribacteroid membrane to mediate molybdate efflux from cytoplasm of plant host cells to the symbiosome. Upon reaching the peribacteroid space, *ModABC* operon would introduce molybdate into the bacteroid. These molybdate could be loaded to the assembly of Fe-Mo cofactor and further for biosynthesis of nitrogenase. PM: plasma membrane; PBM: peribacteroid membrane

## Materials and methods

### Plant materials and growth conditions

Two mutant lines *sen1-1 (Ljsum75*) and *sen1-2* (s88) were produced from ethylmethane sulfonate mutagenesis of ecotype *Gifu* B-129, as reported previously (Kawaguchi et al., 2002). The details of map-based cloning, segregation analysis of F2 progeny from crosses between mutant lines and the WT, and cDNA cloning of these mutant lines have been given in our previous study (Hakoyama et al., 2012). *sen1-1* and *sen1-2* both harbored single nucleotide mutations leading to amino acid substitutions (**Figure S1**).

Seeds of the parent *L. japonicus* ecotype *Gifu* B-129 and the two mutant lines (*sen1-1*, *sen1-2*) were scarified with sandpaper, surface-sterilized with bleach, and then soaked in sterile water and shaken gently at room temperature for 24 h. Subsequently, the seeds were sown on agar (0.8%) plates containing half-strength B&D medium (Broughton and Dilworth, 1971) and left in the dark for 2 days. This was followed by cool-white light illumination (~100 μmol photons m^-2^ s^-1^, 16 h day/8 h night cycle) for 1 week. Nine days after germination the cotyledons were removed from the seedlings to limit Mo supply from cotyledons. The seedlings were then transplanted to a plastic pot filled with vermiculites, where they were watered with B&D solution (Broughton and Dilworth, 1971) with deficient (17 nM) or sufficient (170 nM) Mo. When the plants were transplanted to the pots they were inoculated with the *Mesorhizobium loti* MAFF303099. Ten or 28 days after inoculation, the plants were harvested for nodule morphology observations and Mo concentration determination. The plants were grown in a controlled chamber at 26 °C with 16 h day/8 h night cycle.

### Isolation of bacteroids from the nodule host fractions

Bacteroids were isolated from nodules of the WT and the *sen1-1* and *sen1-2* mutants at 10 or 28 dpi, in accordance with previous studies (Day et al., 1989; Hakoyama et al., 2012). Nodules were homogenized in 50 mM Tris–HCl (pH 7.5) and 0.15M NaCl, and then they were separated into a host plant fraction and a bacteroid fraction by centrifugation. The bacteroids and the nodule host fractions were rinsed three times by ultrapure water and then digested with concentrated HNO_3_ and H_2_O_2_. Concentration of Mo was determined by inductively coupled plasma mass spectroscopy (ICP-MS) (Agilent 7800, Agilent Technologies, USA).

### Heterologous expression of *SEN1* in *S. cerevisiae* and Mo transport activity assay

The ORF of *SEN1* or *sen1-1* was amplified by PCR using cDNA from the nodules and then cloned into the yeast expression vector pYES2 (Invitrogen, Carlsbad, CA), which allows expression of the inserted gene via the galactose-inducible *GAL1* promoter. The *SEN1; pYES2* vector and *sen1-1; pYES2* were introduced into *S.cerevisiae* (BY4741, *MATa his3Δ1 leu2Δ0 met15Δ0 ura3Δ0*) by using the lithium acetate transformation, respectively, and the empty vector pYES2 was used as a negative control. After transformation, the yeast cells were grown on plates with Mo-free selection (SD or SG) (Fred, 2002). To determine Mo uptake activity, yeast cells carrying *SEN1;* pYES2, *sen1-1;* pYES2, or empty vector were cultured to the mid-log phase in Mo-free SD or SG medium and then incubated for 30 min at 30 °C in the same medium containing 170 nM Na_2_MoO_4_. After Mo treatment, the yeast cells were harvested by centrifugation and washed twice in ice-cold ultrapure water. The yeast cells were then re-suspended in 2 mL ice-cold ultrapure water, with the OD_600_ recorded, and were digested with concentrated HNO_3_ and H_2_O_2_ for ICP-MS analysis.

### RNA isolation and qRT-PCR

To investigate the *SEN1* expression levels in different tissues, seedlings were grown in pots containing half-strength B&D solution supplemented with 17 nM, 170 nM, or 1μM Mo. Nodules, roots, and shoots were harvested for RNA extraction 28 days after inoculation with *M. loti* MAFF303099. To investigate the temporal variations in *SEN1* expression in nodules, the nodules samples were harvested at 7, 14, 21, 28, 35, and 42 dpi from plants grown in the half-strength B&D solution supplemented with 170 nM Mo. The RNA extraction, cDNA synthesis, and qRT-PCR were done as described previously (Duan et al., 2017). The primers used for *SEN1* and the internal standard (the *ubiquitin* gene) are shown in **Table S1**.

For the expression analysis of *nifD* and *nifK*, nodules were harvested from WT plants and the *sen1-1* and *sen1-2* mutants at 21 dpi, and bacteroids were prepared from each nodule by centrifugation as described in a previous study (Suganuma et al., 2003). Total RNA was isolated from the bacteroids. The cDNA was then synthesized by reverse-transcriptase (Superscript II, Invitrogen). The primers used for *NifD, NifK* and the internal standard (the *sigA* gene) (Ott et al., 2005), are shown in **Table S1**. Normalized relative expression was calculated by using the ΔΔ*C*t method.

### Immunohistological analysis

The 2079 bp promoter along with the ORF of *SEN1* (there is no intron in the *SEN1* genomic sequence), excluding the stop codon, were amplified by using the primers in **Table S1**. The amplified PCR products were cloned into the *Bam*HI and *Eco*RI sites of pENTR2B vector (Invitrogen). The resulting sequence was subsequently subcloned into pMDC107 vector to create the *pSEN1; SEN1-GFP* constructs by using the Gateway LR reaction (Invitrogen). These constructs were transformed into *Agrobacterium rhizogenes* strain AR1193 for further hairy-root transformation according to the method described in a previous study (Hakoyama et al., 2012). Transgenic hairy roots were inoculated with DsRed-tagged *M. loti* 303099.

Immunostaining was performed according to the methods used by Sauer et al., (2006), with some modification. Briefly, 28-day-old nodules were fixed in 4% (w/v) paraformaldehyde buffered with microtubule-stabilizing buffer (MTSB) (pH 7.4). Then 0.1% Triton X-100 was added, and the mixture was kept overnight at 4 °C. After being washed twice with MTSB and two times with ultrapure water, the fixed samples were embedded in 5% agar in MTSB and sectioned (100-μm) with a vibratome. Afterwards, the sections were placed on microscope slides and incubated with MTSB containing 0.1% (w/v) pectolyase Y-23 (Seishin) at 30 °C for 2 h. They were then reincubated in MTSB containing 0.3% (v/v) Triton X-100 at 30 °C for 2 h. Next, the sections were washed three times with MTSB and blocked with 5% (w/v) bovine serum albumin in MTSB. After the blocking, the sections were treated with primary antibody solution (1: 100 anti-GFP antibody (Thermo Fisher Scientific) in the blocking buffer) overnight at 37 °C and then washed three times. They were then treated with secondary antibody solution (1 : 200 of Alexa Fluor 488 goat anti-rabbit IgG; Molecular Probes, in the blocking buffer) for 2 hours at 37 °C and washed three times. The samples were mounted with 50% (v/v) glycerol in MTSB onto glass slides and observed by confocal laser microscopy (Olympus, FLUOVIEW FV3000). The respective excitation and emission wavelengths were 488 nm and 507-532 nm for the anti-GFP antibody, and 561 nm and 600-650 nm for RFP-tagged rhizobia.

Furthermore, the synthetic peptide CRDMIKSEQGERDLEMAME (positions 111-128 of the SEN1 amino acid sequence) was used to immunize rabbits to obtain antibodies against *SEN1*. The antiserum was purified through a peptide affinity column. Antibody specificity was confirmed by western blot using the total and microsome proteins from 21-dpi nodules. Immunoelectron microscopy for immunolocalization of *SEN1* was conducted according to previous study (Toyooka et al., 2009). Briefly, nodules were collected at 7, 14, 21 dpi and were frozen in a high-pressure freezer (EM-ICE; Leica, Vienna, Austria) by using the fixation solution (0.25% (w/v) glutaraldehyde and 0.1% uranyl acetate (w/v) in acetone). The nodule samples were then replaced in methanol for 5 min each at 4 °C and in LR White hard-type resin at 3:1, 1:1 and 1:3 for 1 h each; they were then placed in 100% LR White resin twice for 1 h each. The samples were finally embedded in gelatin capsules in 100% LR White resin at −20 °C for 72 h under ultra-violet light. Ultra-sections were cut with a diamond knife (Ultra: Diatome, Nidau, Switzerland) and a Leica EM UC-7 ultramicrotome (Leica Microsystems, Vienna, Austria). The sections on nickel grids were blocked with a blocking buffer (Block Ace Powder; DS Pharma Biomedical Co., Ltd) for 30 min. After being blocked, the sections were labeled with anti-SEN1 antibody (1: 20) for 5 h and anti-rabbit goat IgG antibody (1:20) conjugated with 12-nm colloidal gold particles diluted in blocking buffer. The sections were stained with 4% uranyl acetate (w/v) for 10 min and visualized under a transmission electron microscope (JEOL JEM-1400, Jeol, USA).

## Supplemental materials

**Supplemental Figure 1.**Predicted tertiary structure model of SEN1 protein using AlphaFold v3 and Schematic domain structure of the SEN1 protein and the point-mutation positions of the two *sen1* mutants.

**Supplemental Figure 2.**Mo concentration in the roots (A, C) and shoots (B, D) of WT and mutant lines grown under sufficient or deficient Mo supply.

**Supplemental Figure 3.**The specificity of antibody against SEN1 protein by western blot in the crude total protein and microsome protein of nodules from WT.

**Supplemental Figure 4.**Subcellular localization of SEN1 in the infected cells of nodules at 7 dpi, 14 dpi and 21 dpi, respectively, using anti-SEN1 antibody.

**Supplemental Figure 5.**The frequency of colloidal gold particles observed in PBM, nodule host cells (intracellular components excluding symbiosome), and bacteroids in the cross-sections of nodule samples harvested from different dpi.

**Supplemental Table 1.**The primers used in this study.

## Acknowledgements

This work was supported by the Grant-in-Aid for Scientific Research to T. F. (No. 20F20098, 18H05490 and 19H05637), and by a Fellowship from the Japan Society for the Promotion of Science (JSPS) to Q. C. We thank Dr. Marcel Beier for the help during the experiment of western blot and immunostaining.

## Author contributions

T.H., M.H. and T.F conceive the research plans; Q.C, T.H., M.H., and T.F. designed major aspects of the project; Q.C, T.H., K.T. and M.S. performed the experiments; Q.C, T.H., K.T. and M.S. analyzed the data; Q.C. and T.F. conceived the project and wrote the article with contributions of all the authors; T.F. supervised and finalized the manuscript. T.F. agrees to serve as the author responsible for contact and ensures communication.

